# Coupling of metataxonomics and culturing improves bacterial diversity characterization and identifies a novel *Rhizorhapis* sp. with metal resistance potential in a multi-contaminated waste sediment

**DOI:** 10.1101/2022.02.20.481181

**Authors:** José A. Siles, Andrew J. Hendrickson, Norman Terry

## Abstract

Long-term contaminated environments have been recognized as potential hotspots for bacterial discovery in taxonomic and functional terms for bioremediation purposes. Here, bacterial diversity in waste sediment collected from a former industrial dumpsite and contaminated with petroleum hydrocarbon and heavy metals was investigated through the parallel application of culture-independent (16S rRNA gene amplicon sequencing) and -dependent (plate culturing followed by colony picking and identification of isolates by 16S rRNA gene Sanger sequencing) approaches. The bacterial diversities retrieved by both approaches greatly differed. *Bacteroidetes* and *Proteobacteria* were dominant in the culture-independent community, while *Firmicutes* and *Actinobacteria* were the main culturable groups. Only 2.7% of OTUs (operational taxonomic units) in the culture-independent dataset were cultured. Most of the culturable OTUs were absent or in very low abundances in the culture-independent dataset, revealing that culturing is a useful tool to study the rare bacterial biosphere. One culturable OTUs (comprising only the isolate SPR117) was identified as a potential new species in the genus *Rhizorhapis* (class *Alphaproteobacteria*) and was selected for further characterization. Phytopathogenicity tests showed that *Rhizorhapis* sp. strain SPR117 (ATCC TSD-228) is not pathogenic for lettuce, despite the only described species within this genus, *Rhizorhapis suberifaciens*, is causal agent of the lettuce corky root disease. The genome of the strain SPR117 was sequenced, assembled in 256 contigs, with a length of 4,419,522 bp and a GC content of 59.9%), and its further annotation revealed the presence of genes related to the resistance to arsenic, copper, iron, and mercury, among other metals. Therefore, the coupling of metataxonomics and culturing is a useful tool to obtain not only an improved description of bacterial communities in contaminated environments, but also to isolate microorganisms with bioremediation potential.

## 1. Introduction

Pollutants such as heavy metals, total petroleum hydrocarbons (TPHs), and polycyclic aromatic hydrocarbons (PAHs) are continuously released into the environment as a consequence of industrial activities, their associated disposal operations, and accidental spills (Brassington et al., 2007; Chen et al., 2015; Liu et al., 2018). Most of them are not easily decomposable or not degradable at all and accumulate in terrestrial environments, exerting a negative impact on quality services of ecosystems, animal life, and human health (Okereafor et al., 2020). Continuous and long-term exposure to contaminants usually leads to perturbations in bacterial communities (Feng et al., 2018). These alterations are usually driven by selective pressure, which favors those bacterial populations with the ability of tolerating the introduced contaminants and/or using them to their advantage (Lukhele et al., 2021; Thavamani et al., 2012; Valls and de Lorenzo, 2002). Mobile genetic elements have shown to have a critical role in this process (Dziewit et al., 2015). The study of bacterial communities in ecosystems contaminated with hydrocarbons and/or metals is thus of interest since they may be a valuable reservoir for bacterial discover in functional and taxonomic terms (Bao et al., 2017). In this context, with the introduction of high-throughput sequencing techniques, microbial metataxonomic and metagenomic studies in hydrocarbon- and metal-polluted sites have proliferated in detriment of culture-dependent surveys. This has shed light on the ecology and functionality of bacterial communities in these environments. For example, the “gamma-shift” is now a well-known phenomenon consisting in the enrichment of soil microbial communities in *Gammaproteobacteria* populations followed TPH entrance into the ecosystem (Dong et al., 2015; Militon et al., 2010; Siles and Margesin, 2018). Another example is the metagenomic work of Delmont et al. (2015), which showed that metals exert a positive selective pressure on genes related to their microbial resistance and reconstructed multiple bacterial genomes directly from metal-contaminated soils.

Nonetheless, in the last few years, as genomic sequencing data accumulate, there has been a new interest in culturing to obtain as-yet-uncultured bacteria since isolates are still required to fully discover the physiology and ecology of bacteria and improve reference databases for sequence annotation (Lewis et al., 2020; Thrash, 2019, 2021). In fact, most of the genes recovered from bacterial genomes or metagenomes cannot currently be annotated, in part, because our ability to link DNA sequences to gene products is mainly based on the study of only a few bacterial isolates (Carini, 2019). This renewed motivation to culture bacteria from terrestrial ecosystems is also based on recent advances in culturing techniques and the enormous amount of molecular data probing the limitless microbial diversity in these environments, which we know very little about (Carini, 2019; Delgado-Baquerizo, 2019; Delgado-Baquerizo et al., 2018; Pham and Kim, 2012). Moreover, although the classical “1% culturability paradigm” (which means that around 99% of environmental microorganisms are unculturable) is no longer accurate, most bacterial taxa remain uncultured (Martiny, 2019; Steen et al., 2019). In the last few years, new culturing approaches have been developed (Lewis et al., 2020), which along with the traditional plate-based methods have led to the isolation of new bacterial taxa that have aroused considerable interest because of the novelty of the isolated microbes or because the cultured microorganisms provided an improved understanding of certain natural processes (Lewis et al., 2020). In this context, traditional plate culturing, with subsequent colony picking, has still been shown to be a valid and successful method for the culture-dependent characterization of terrestrial bacterial communities and the isolation of “uncultured” bacteria (Nguyen et al., 2018; Thrash, 2021). The use of (i) culture media poor in nutrients (oligotrophic), extracted from the environment (e.g., soil) or supplemented with specific nutrients or chemical requirements for bacterial growth; (ii) coculture with helper bacteria; (iii) long incubation periods (in the range of up to several months); (iv) alternating incubation temperatures; and (v) methods to physically reduce the number and diversity of bacteria in the sample to be analysed (e.g., filtration, density-gradient centrifugation or extinction-dilution) are helpful strategies to isolate novel bacterial strains through plate-based methods (Nguyen et al., 2018; Vartoukian et al., 2010). Diffusion chambers (e.g., i(isolation)Chip) have also successfully been applied to culture novel bacterial taxa (Kaeberlein et al., 2002; Thrash, 2021).

A new approach in the study of bacterial communities in terrestrial ecosystems comprises the coupling of 16S rRNA gene amplicon sequencing and culturing (Hamood Altowayti et al., 2020; Spini et al., 2018; Stefani et al., 2015). This provides insights into the taxa that culturing is biased towards and reveals whether the isolated strains belong to the abundant or rare bacterial biosphere (Pascual et al., 2016; Shade et al., 2012). Bacterial communities in environments such as cropland (Lee et al., 2016; Shade et al., 2012) and forest (VanInsberghe et al., 2013) soils have been investigated through the coupling of metataxonomics and culturing. However, to the best of our knowledge, this approach has not yet been applied to long-term industrial waste dumpsites, which have been identified as hotspots for discovery of novel bacteria with bioremediation abilities (Muhonja et al., 2018; Singh et al., 2015). This represents a research gap, which we aimed to address in the current study. The objective of the present work was to characterize the bacterial diversity from waste sediment collected from a former industrial dumpsite (located in Bay Point, CA, USA) and contaminated with TPHs, PAHs, and metals by applying culture-independent (16S rRNA gene amplicon sequencing) and -dependent (plate culturing with further colony picking and identification of isolates by 16S rRNA gene amplification and Sanger sequencing) approaches. We hypothesized that bacterial culturable diversity would be greatly different and would represent only a small fraction of that retrieved through amplicon sequencing. We also hypothesized that the studied sediment would be rich in potential novel species of bacteria due to its peculiar characteristics, which would be revealed by metataxonomics by yielding a high proportion of unclassified sequences at the different taxonomic levels. Since one of the isolates was a potential new species within the genus *Rhizorhapis* -with important agronomic and environmental implications-, we decided to gain more insight into the features of this strain through genome sequencing.

## 2. Materials and methods

### 2.1. Site description and sampling

The studied site is located along the waterfront next to Suisun and Honker Bays in Bay Point, California, USA (geographical coordinates: 38°02’12.7” N, 121°56’52.9” W) and is a former solid waste management unit (SWMU). Its dimensions are roughly 800 m (north to south) by 370 m (east to west), with a slightly trapezoidal shape. The SWMU (which is currently owned by The Pacific Gas and Electric (PG&E) Company) was constructed in the late 1940s to receive stormwater and wastewater discharges from commercial production plants and was active until 1980. Currently, the SWMU is composed of a layer of waste sediment (whose thickness ranges from 7.5 cm to a maximum of 150 cm) above natural peat and younger bay mud. The waste sediment is comprised of a varying mixture of residues from the former manufacturing plants, carbon black (fly ash residue from coal burning industry, particle size <10 μm), and organic matter (CH2M-Hill, 2010). The sediment is black, highly saline (due to the influence of marine waters), and presents contamination by TPHs, PAHs, and metals (Khan et al., 2019; Xia et al., 2017). This site represents a serious environmental concern and will undergo a process of bioremediation in the near future. At the time of this study, the SWMU was partially covered by water to mitigate the odors that emanate from the sediment and to prevent the release of sediment dust. In the areas without water cap, SWMU was irregularly colonized by saltgrass (*Distichlis spicata*), pickleweed (*Salicornia virginica*), and sea purslane (*Sesuvium verrucosum*) (Xia et al., 2017). The climate in the area is typically Mediterranean, with a mean annual temperature of 15.4 °C and annual precipitation of approximately 339 mm.

For sampling, an area of approximately 200 m2, free of water cap, was delimited and three equidistant sampling spots were identified and selected to collect three replicate sediment samples (one from each spot). Each sample was a composite of five subsamples from the top 20 cm of the layer sediment: four subsamples orthogonally collected in a 2-m radius from a central subsample. Sampling points were free of vegetation. In total, around 500 g of sediment were collected per sample. Then, samples were transported to the lab in cooled boxes and subsequently homogenized, sieved (2 mm mesh), and stored at 4 °C for physicochemical characterization and bacterial plate culturing or at −80 °C for DNA extraction. Sampling and processing of the samples were carried out using sterilized tools to avoid contamination.

### 2.2. Physicochemical characterization of the waste sediment

Sediment samples were physicochemically characterized (texture, pH, electrical conductivity, total and organic carbon I, total nitrogen (N), and phosphorus) following standard methods at the UC Davis Analytical Lab (Davis, CA, USA). Levels of TPHs in the waste sediment were investigated by firstly extracting them through ultrasonic extraction following the EPA 3550C method and by further injecting an aliquot of the extract into a gas chromatograph equipped with a flame ionization detector according to the EPA 8015B method. PAHs were quantified by firstly extracting samples through pressurized fluid extraction according to the EPA 3545 method and by subsequently characterizing extracts by gas chromatography-mass spectroscopy following the EPA 8270D method. The EPA 6020 method was used to quantify the levels of heavy metals in the sediment samples through the application of inductively coupled plasma mass spectrometry. The analyses were done at Eurofins Calscience (Garden Grove, CA, USA).

### 2.3. Culture-independent bacterial diversity

Culture-independent bacterial diversity in the waste sediment was assessed through Illumina 16S rRNA gene amplicon sequencing (also known as 16S rRNA gene metabarcoding or metataxonomics). To do that, firstly, total DNA from each of the three sediment samples was extracted in triplicate from 250 mg of sediment fresh mass using Dneasy PowerSoil Kit (Qiagen, Valencia, CA, USA) following the manufacturer’s instructions. Triplicate DNA extracts from each sample were then pooled and DNAs were purified using DNA Clean & Concentrator Kit (Zymo Research, Irvine, CA, USA) according to manufacturer’s instructions. The quality of DNAs was spectrophotometrically checked by NanoDrop (Thermo Fisher Scientific, Waltham, MA, USA) based on the absorbance ratios A260/A280 and A260/A230. Subsequently, DNAs were quantified using QuantiFluor™ dsDNA System (Promega, Madison, WI, USA) and DNA concentrations were standardized.

A fragment of the 16S rRNA gene was amplified using the primers 27F (5′- AGRGTTTGATCMTGGCTCA-3′) and 519R (5′- CCCCGYCAATTCMTTTRAGT -3′). These primers were selected since they capture the V1-V3 hypervariable regions of the bacterial 16S rRNA gene, which have been shown to provide higher taxonomic resolution for lower rank taxa than other 16S hypervariable regions (Bukin et al., 2019). PCR reactions were conducted using the HotStarTaq *Plus* Master Mix Kit (Qiagen, Valencia, CA, USA) and barcoded forward primers, under the following thermal conditions: (i) 94°C for 3 min; (ii) 30 cycles of 94°C for 30 s, 53°C for 40 s, and 72°C for 60s; with (iii) a final elongation step at 72°C for 5 min. The success of the amplifications was checked in a 2% agarose gel and PCR products were subsequently purified using Agencourt AMPure XP magnetic beads kit (Beckman Coulter, Brea, CA, USA), quantified with the QuantiFluor™ dsDNA System (Promega, Madison, WI, USA), and pooled in equal proportions. The pooled product was then used to prepare the Illumina DNA library. Paired-end (PE) sequencing (2 × 300) was performed on an Illumina MiSeq sequencing platform (Illumina, San Diego, CA, USA) at MR DNA (www.mrdnalab.com, Shallowater, TX, USA).

Bioinformatic processing of the sequences was conducted using the USEARCH pipeline and UPARSE-OTU algorithm (Edgar, 2013). Briefly, PE sequences were firstly merged with the command - *fastq_mergepairs.* Then, reads were quality-filtered allowing a maximum e-value of 1.0, trimmed to 450-bp (base pair), dereplicated, and sorted by abundance (removing singleton sequences), prior chimera detection and OTU (operational taxonomic unit) determination at 97% sequence identity. Finally, original trimmed and high-quality sequences were mapped to OTUs at the 97% identity threshold to obtain one OTU table. The taxonomic affiliation of each OTU was obtained using RDP (ribosomal database project) taxonomic classifier (Wang et al., 2007) against 16S rRNA training set 18 with a 80% confidence threshold. OTUs with average (*n*=3) relative abundances above 1% were considered as abundant “taxa”, those with relative abundances below 0.1% were defined as rare and those with relative abundances between 0.1% and 1% were identified as “intermediate” OTUs (Fuhrman, 2009).

### 2.4. Culture-dependent bacterial diversity

Culturable heterotrophic bacteria inhabiting the waste sediment were studied by plate culturing (using two culture media differing in their nutrient contents), followed by colony picking and identification of isolates by 16S rRNA gene amplification and Sanger sequencing. To do this, one suspension per sample (*n*=3) was prepared by shaking 10 g of sediment with 90 ml of sterile saline solution (0.85% NaCl, w/v) for 30 min. Appropriate dilutions of soil suspensions were surface spread onto LB (Lysogeny Broth, Merck Millipore, Burlington, VT, USA) and R2A (Reasoner’s 2A; Research Products International, Mt Prospect, IL, USA) agar plates, with a pH of 8.2, and containing cycloheximide (200 μg mL−1), to exclude fungal growth. While LB is a rich medium, R2A is a relatively nutrient-poor medium that favors the isolation of oligotrophic bacteria (Molina-Menor et al., 2021). The plates were wrapped in plastic bags to prevent evaporation and incubated during 50 days at 20 °C. Afterwards, using the plates with the least number of colonies, 80 colonies growing on LB plates and 70 colonies growing on R2A plates (for a total of 150 colonies) were transferred to new Petri dishes and incubated for 2 weeks in order to identify unpurified and non-growing colonies. Finally, a total of 140 isolates −75 of them isolated from LB medium and 65 from R2A-were selected for taxonomic identification by 16S rRNA gene sequence analysis. To do this, genomic DNA from each of the isolates was extracted using the Dneasy UltraClean Microbial Kit (Qiagen, Valencia, CA, USA) following the manufacturer’s instructions. The primers fD1 (5’- AGAGTTTGATCCTGGCTCAG - 3’) and rP2 (5’-ACGGCTACCTTGTTACGACTT-3’) were used for almost complete 16S rRNA gene amplification (Weisburg et al., 1991). PCR reactions were carried out in a final volume of 50 μL with 25 μL Taq 2X Master Mix (New England BioLabs, Ipswich, MA USA), 1 μL each forward and reverse primers (10 μM) (Genewiz, South Plainfield, NJ, USA), 1 μL DNA solution, and 23 μL H_2_O. The thermal cycling program was as follows: (i) 95°C for 5 min; (ii) 35 cycles of 95 °C for 30 s, 53°C for 30 s, and 72°C for 60s; and (iii) a final elongation step of 72°C for 5 min. 16S rRNA gene PCR products were visualized on an agarose gel stained with SYBR™ safe (Thermo Fisher Scientific, Waltham, MA, USA), purified using GENEJET PCR purification kit (Thermo Fisher Scientific, Waltham, MA, USA), and Sanger sequenced (UC Berkeley DNA Sequencing Facility; Berkeley, CA, USA).

The sequences were manually edited using the software MEGA ver. X (Kumar et al., 2018) and the nearest phylogenetic neighbors were determined for each strain using the EzTazon-e Database (Kim et al., 2012). The information obtained from the taxonomic affiliation of each isolate was used to describe the culturable bacterial diversity in the waste sediment. Additionally, the 16S rRNA gene sequences of the isolates were clustered into OTUs and phylogenetically related to the OTUs obtained through 16S rRNA gene metabarcoding at a 97% identity threshold using the program VSEARCH (Rognes et al., 2016) and distance-based greedy clustering.

### 2.5. Phylogenetic analysis and genome sequence of *Rhizorhapis* sp. strain SPR117 (ATCC TSD-228)

Almost complete 16S rRNA gene sequence of *Rhizorhapis* sp. strain SPR117 was deposited at DDBJ/EMBL/GenBank under the accession number OM362825. Phylogenetic analysis of SPR117 16S rRNA gene sequence and those of other related strains was performed using the software MEGA ver. X. Additional 16S rRNA gene sequences of related strains were retrieved from public databases. Sequences were firstly aligned with the Clustal W software (Thompson et al., 1994) and the phylogenetic tree was then reconstructed by the neighbour-joining algorithm (Saitou and Nei, 1987). The genetic distances were calculated by using Kimura’s two-parameter model (Nishimaki and Sato, 2019) and the complete deletion option. The resultant tree topologies were evaluated by bootstrap analysis based on 1000 replicates.

For whole-genome sequencing, *Rhizorhapis sp.* strain SPR117 was grown during 7 days in R2A broth at 20 °C and its genomic DNA (gDNA) was extracted using the Dneasy UltraClean Microbial Kit (Qiagen, Valencia, CA, USA) according to the manufacturer’s instructions. GDNA was then quality-checked, quantified, and used for the preparation of a NEBNext® Ultra™ DNA library (New England Biolabs, Ipswich, MA, USA) following the manufacturer’s recommendations. The DNA library was loaded on an Illumina HiSeq (Illumina, San Diego, CA USA) instrument and sequenced using a 2×150 PE configuration in High Output Run mode at Genewiz (South Plainfield, NJ, USA).

Raw reads were quality-checked using FastQC v0.11.5 and filtered using Trimmomatic v.0.38 (ILLUMINACLIP = TRUE; LEADING = 3; TRAILING = 3; SLIDINGWINDOW = 4:30; MINLEN = 90). Filtered PE sequences were then de novo assembled with SPAdes v. 3.13.0 (Bankevich et al. (2012); -no trim, 2 racon iterations, 2 pilon iterations, 5 minimum contig coverage and 300 bp minimum contig length-) integrated in PATRIC Bioinformatics Resource Center (Davis et al., 2020). Subsequently, contigs were annotated using the RAST (Rapid Annotations using Subsystems Technology, Brettin et al. (2015)), Prokka v.1.14.6 (Seemann, 2014), and the NCBI Prokaryotic Genome Annotation Pipeline (Tatusova et al., 2016). CheckM algorithm was used for assessing the genome quality (Bankevich et al., 2012).

### 2.6. Phytopathogenicity assay

The strain SPR117 was tested for pathogenicity to two different lettuce varieties: Waldmann’s Dark Green (*Lactuca sativa* L. cv. Waldmann’s dark green) and Romaine (*Lactuca sativa* L. var. *longifolia*). To do this, seeds of the two lettuce varieties were surface disinfected by immersion in 2% (v/v) hydrogen peroxide for 5 min. Seed germination was carried out at 25 °C in trays containing sterilized vermiculite as substrate for one week. Afterwards, a 7-day old lettuce plant was planted in an 8-cm diameter pot containing sterilized potting mix substrate. In total, 24 pots were prepared for each lettuce variety. Three days after planting, and once a week afterwards, 12 of the 24 pots of each lettuce variety were inoculated with 5 mL of a cell suspension of SPR117 with an optical density of 0.380, equivalent to a concentration of 1.4 × 107 colony forming units mL^−1^. The SPR117 inoculum was prepared by growing the strain during 7 days in R2A broth at 20 °C under shaking conditions, centrifuging to remove culture medium, and resuspending the pellet in sterile saline solution (0.85% NaCl, w/v). The 12 control pots received 5 mL of sterile saline solution. Lettuce plants were watered as needed with sterilized deionized water. The experiment was conducted in a greenhouse with controlled light and humidity conditions. Four weeks after the first bacterial inoculation, the plants were evaluated for lettuce corky root severity based on a 0–9 scale according to Brown and Michelmore (1988) and dry weight of shoot and root.

### 2.7. Statistical analyses

Non-parametric univariate PERMANOVA (Permutational Analysis of Variance) was used to check whether root and shoot dry weight of lettuce plants was significantly affected by the inoculation of *Rhizorhapis* sp. strain SPR117. For this analysis, the *adonis* function in the R package *vegan* was used (Oksanen et al., 2013). Data visualizations were performed using the R package *ggplot2* (Wickham, 2016) and CorelDRAW ver. 2020.

### 2.8. Strain and data accessibility

*Rhizorhapis* sp. strain SPR117 was deposited at the American Type Culture Collection (ATCC) under the strain designation ATCC TSD-228.

The draft-genome sequence of *Rhizorhapis* sp. strain SPR117 was deposited at DDBJ/EMBL/GenBank under the accession number JAKKIE000000000 (BioProject no. PRJNA800004 and BioSample no. SAMN25221871). The version described in this paper is the first version, JAKKIE000000000. Raw genome sequences of *Rhizorhapis* sp. SPR117 were deposited in the GenBank SRA database under accession number SRP361685.

The raw sequences of the 16S rRNA gene amplicon sequencing analysis were deposited in the SRA database under BioProject accession number PRJNA800331.

## 3. Results and discussion

### 3.1. Physicochemical properties of the waste sediment

The sediment was characterized by presenting an extremely high C/N ratio (199), alkaline pH (8.2), high salinity (electrical conductivity, 1.2 mS cm^−1^), and high levels of contamination by TPHs (20.1 g kg^−1^ sediment), PAHs (370 mg kg^−1^ sediment; predominating: benzo(*ghi*)perylene, pyrene, indeno(1,2,3-*cd*)pyrene, benzo(*a*)pyrene, fluoranthene), and heavy metals (chromium, barium, zinc, copper, lead and nickel) (Table S1). These data confirmed the peculiar physicochemical characteristics of the studied waste sediment.

### 3.2. Culture-independent and -dependent bacterial diversity in the waste sediment

The metabarcoding assay yielded 98,948 PE high-quality 16S rRNA gene sequences, which were distributed among 625 OTUs. Of these OTUs, 621 were classified at kingdom level as *Bacteria* (using an 80% confidence threshold) and the remaining 4 as *Archaea*. Only bacterial OTUs were retained for further taxonomic analyses. Culture-independent bacterial diversity was distributed among 20 different phyla, being the predominant ones *Bacteroidetes* (the average percentage of classified sequences in this phylum was 52.4), *Proteobacteria* (14.2%), *Actinobacteria* (7.6%), *Planctomycetes* (4.5%), *Saccharibacteria* (3.3%), *Verrucomicrobia* (3.0%), and *Balneolaeota* (2.7%); these six phyla accounted for more than 87% of total sequences (Fig. 1a). *Flavobacteriia* class predominated among *Bacteroidetes*, *Gammaproteobacteria* among *Proteobacteria*, *Planctomycetacia* among *Plantomycetes,* and *Acidimicrobiia* among *Actinobacteria* (Fig. 1b). These findings concur with those of previous works showing that *Bacteroidetes*, *Proteobacteria*, and *Actinobacteria* usually predominate in long-term contaminated soils with metals and hydrocarbons (Khan et al., 2019; Pan et al., 2020; Siles and Margesin, 2018). At OTU level, 11 OTUs were regarded as abundant (i.e., with relative abundances >1%) and belonged to the phyla *Actinobacteria, Bacteroidetes*, *Chloroflexi*, *Proteobacteria*, *Saccharibacteria*, and *Verrucomicrobia* (Table S2). Among these OTUs, the Otu1, with a relative abundance of 47%, clearly predominated and was taxonomically affiliated to the genus *Hoppeia* (although only with a confidence threshold of 56%, Table S2). The only species described in this genus (namely, *Hoppeia youngheungensis*) was isolated from tidal flat sediment collected from an island in the West Sea of Korea (Kwon et al., 2014), which evidences that marine environments are common habitats for members of this genus. The great influence of marine environment on the studied site would thus explain why the bacterial diversity harbored by the waste sediment comprises the genus *Hoppeia.*

**Fig. 1.**
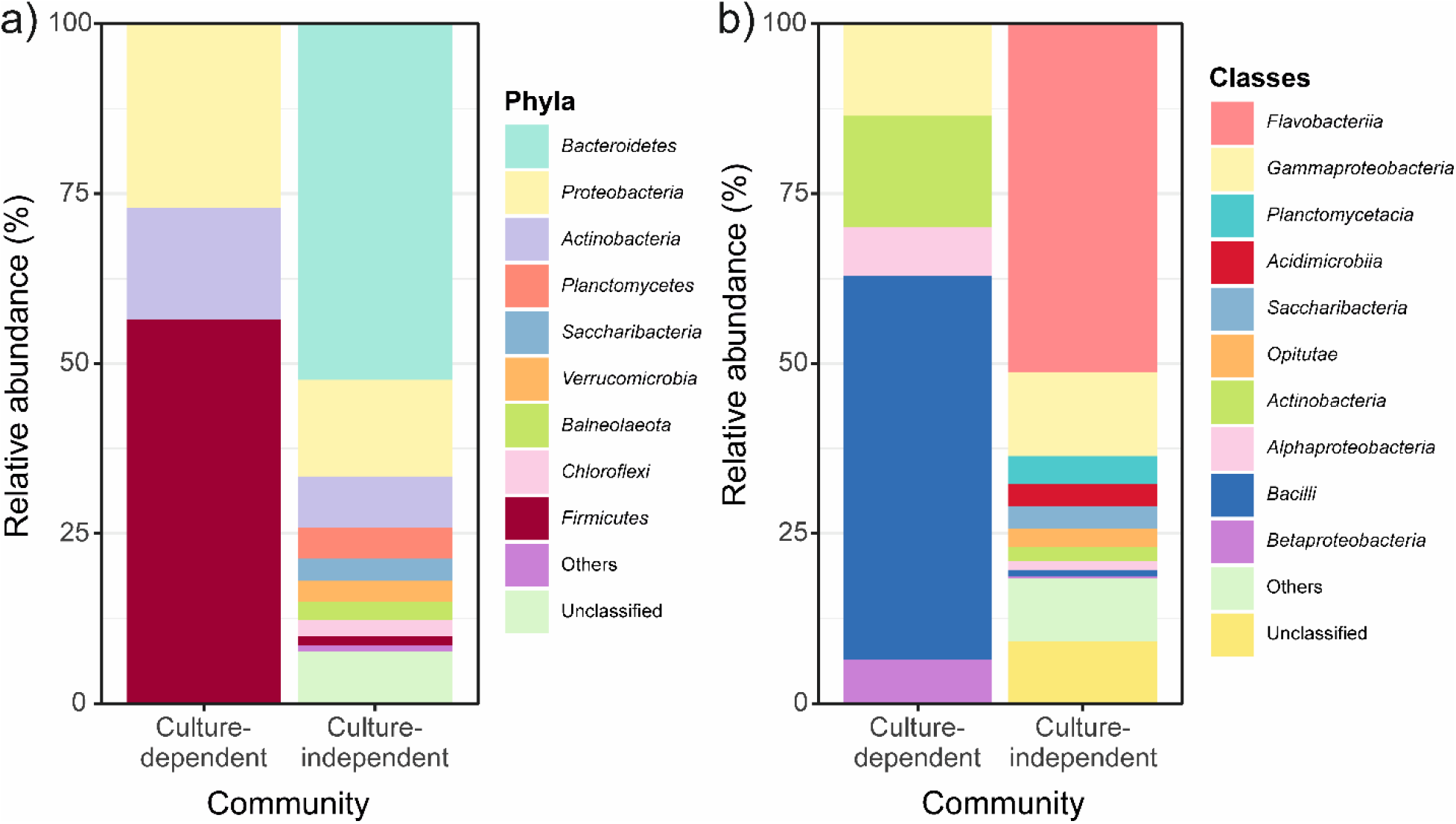
Relative abundance of the phyla (a) and classes (b) conforming culture-dependent (culturing) and - independent (16S rRNA gene amplicon sequencing) bacterial communities in the studied waste sediment.

76.1% of sequences could not be classified at genus level. This result can have two possible explanations: (i) the sequence length was too short for accurate classification or (ii) the reference database was not complete enough and some comparative elements were missing for the classification of all the bacterial sequences (Rachid et al., 2013). In the present study, since (i) a much lower proportion of unclassified sequences at different taxonomic levels was obtained in previous works using the same approaches as those utilized here for taxonomic affiliation of reads with a similar length (Margesin et al., 2017) and (ii) most of the sequences were successfully classified at kingdom (*Bacteria*) and phylum levels, we explain the high proportion of sequences without an effective taxonomic classification as a consequence of the presence of a high proportion of undescribed bacterial diversity in the studied sediment (Delgado-Baquerizo, 2019). In light of these results, we presumed that bacterial culturing would result in the isolation of a high proportion of as-yet-uncultured taxa.

The culture-dependent characterization of the bacterial community in the waste sediment resulted in the isolation and successful identification of 140 strains. The culturable bacterial community was dominated by *Firmicutes* (56.4%), although isolates belonging to the phyla *Proteobacteria* (27.1%) and *Actinobacteria* (16.4%) were also detected (Fig. 1a). Despite we (i) used both rich and oligotrophic culture media for bacterial isolation, (ii) adjusted the pH of media to that of the sediment (i.e., 8.2), and (iii) used a rather long incubation time, the culturable diversity differed from that recovered through the 16S metabarcoding assay, as we initially hypothesized. In this way, only 2.3% of the OTUs in the metabarcoding library were cultured (Fig. 2 and Table S3). These dramatic differences between culture-dependent and - independent bacterial diversities have previously been reported (Lee et al., 2016; Siles et al., 2014) and are a consequence of our incapacity to reproduce in the laboratory the particular abiotic conditions (nutrient concentrations and forms, temperature, humidity, oxygen -or other gases-levels, light, etc.) that most bacteria need to growth (Hugenholtz, 2002; Molina-Menor et al., 2021). Furthermore, the growth of specific taxa depends on interactions between species and metabolic cooperation, which are largely unknown and really hard to mimic under experimental conditions (Molina-Menor et al., 2021). In this way, until recently, it has been believed that culturing only offers a biased and useless description of the bacterial communities of a given environment (Ritz, 2007). However, an increasing number of works are demonstrating that culturing and metataxonomics are complementary to fully describe the microbial diversity in a given environment (Shade et al., 2012; Stefani et al., 2015; Youseif et al., 2021). In our study, phylogenetic comparison of the culture-dependent and -independent diversities at OTU level showed that a total of 21 OTUs were detected only by culturing (Fig. 2). Previous works dealing with this topic have found similar patterns. For example, Lee et al. (2016) found that 73 OTUs were detected only by cultivation after studying the bacterial diversity of tomato rhizosphere through the parallel application of culturing and 16S metabarcoding. Shade et al. (2012) reported that up to 601 OTUs were uniquely retrieved by culture-dependent approaches during the investigation of the bacterial community in an agricultural soil. It has been argued that bacterial taxa recovered only by culturing are rare members of the microbial community, which are detected only by this approach because sequencing depth was insufficient (Hiergeist et al., 2015; Lee et al., 2016). We observed that most of the cultured OTUs in the metabarcoding dataset represented rare members of the culture-independent community (Table S3). Likewise, none of the abundant OTUs in the metabarcoding dataset was cultured (Table S2 and Table S3). Therefore, culturing seems to be especially useful to dig deeper into the rare biosphere of the studied waste sediment. Members of the rare biosphere are essential for the maintenance of microbial alpha and beta diversity and represent a seed bank of genetic and metabolic diversity that responds to different biotic or abiotic changes, contributing to the resilience of the community (Lasa et al., 2019; Lynch and Neufeld, 2015; Saw, 2021).

**Fig. 2.**
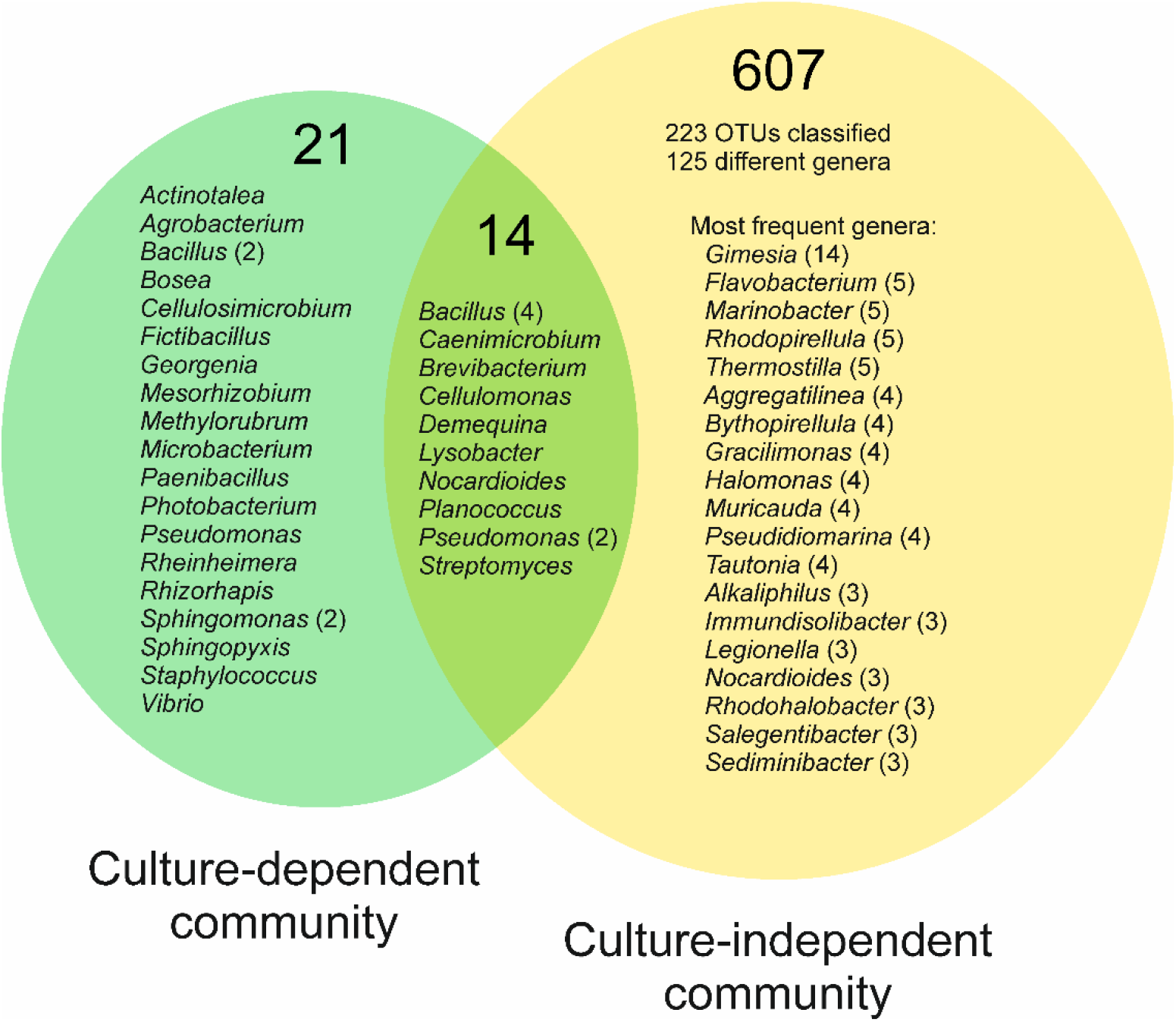
Venn diagram showing the number and taxonomic affiliation (at genus level) of unique and shared OTUs in culture-dependent (culturing) and -independent (16S rRNA gene amplicon sequencing) bacterial communities from the studied waste sediment. Numbers in parenthesis denote the numbers of times that a specific genus was found.

Approximately 79% of the strains shared ≥98.7% sequence identity with the closest known species in the EzTaxon Database, while ca. 21% of the isolates (sharing identities <98.7%) could constitute potential novel species (Table S3). These results suggest that plate culturing is still a suitable method for the isolation of as-yet-uncultured bacteria in terrestrial ecosystems, especially when long incubation periods and other strategies are considered (Thrash, 2021; Vartoukian et al., 2010). For example, 79 different isolates of *Planctomycetes* were recently cultured for the first time using selective enrichment, antibiotic treatment and solid media streaking combined with colony picking (Wiegand et al., 2019). However, it is well known that untargeted cultivation requires substantial amounts of time and resources to be successful. New approaches in culturing tend thus to be more targeted from a functional or taxonomic perspective (Lewis et al., 2020). In this way, lists of most-wanted microbes are being created (Carini, 2019). Reasons to include a specific microorganism in these lists are that it (i) belongs to a poorly characterized group, (ii) is key for a specific process, biogeochemical cycle, or bioremediation, and/or (iii) can potentially produce natural products (Carini, 2019; Lewis et al., 2020). Once one specific microorganism has been targeted for its isolation, particular cultivation strategies can be designed according to its lifestyle. Innovations to cultivate new bacterial taxa include (i) membrane diffusion-based cultivation methods (e.g., iChip or soil substrate membrane system), (ii) microfluidics-based cultivation approaches (e.g., SlipChip), (iii) cell sorting-based techniques (e.g., Raman-activated cell sorting), (iv) fluorescence in situ hybridization of live cells, and (v) reverse genomics (Lewis et al., 2020).

Among all the potential novel strains discovered in the present work, the isolate SPR117 (which was isolated from the medium R2A and comprised the singleton Otu31, Table S3) captured our attention since its closest known strain was, with a pairwise sequence similarity of only 96.88%, *Rhizorhapis suberifaciens* CA1^T^. This finding suggests that the isolate SPR117 represents a new species in the genus *Rhizorhapis*. This assumption was supported by a neighbour-joining phylogenetic tree analysis, which showed that the strain SPR117 -forming a single tree branch- was closely related, but clearly distinct, from a cluster grouping different strains of *R. suberifaciens*. Unfortunately, these branches were not supported by high bootstrap values (Fig 3). *Rhizorhapis* cluster was closer related to the *Sphingobium* cluster than the *Sphingomonas* one (Fig. 3). On the other hand, the OTU containing the isolate SPR117 was not phylogenetically related to any of the OTUs in the amplicon sequencing dataset (Table S3). We thus assume that the isolate SPR117 is a rare member of the bacterial community in the waste sediment, which was not captured by the 16S amplicon sequencing assay. In light of the agronomic (Francis et al., 2014) and environmental importance (Brereton et al., 2020) of the genus *Rhizorhapis*, we decided to gain more insights into the characteristics of the strain SPR177.

**Fig. 3.**
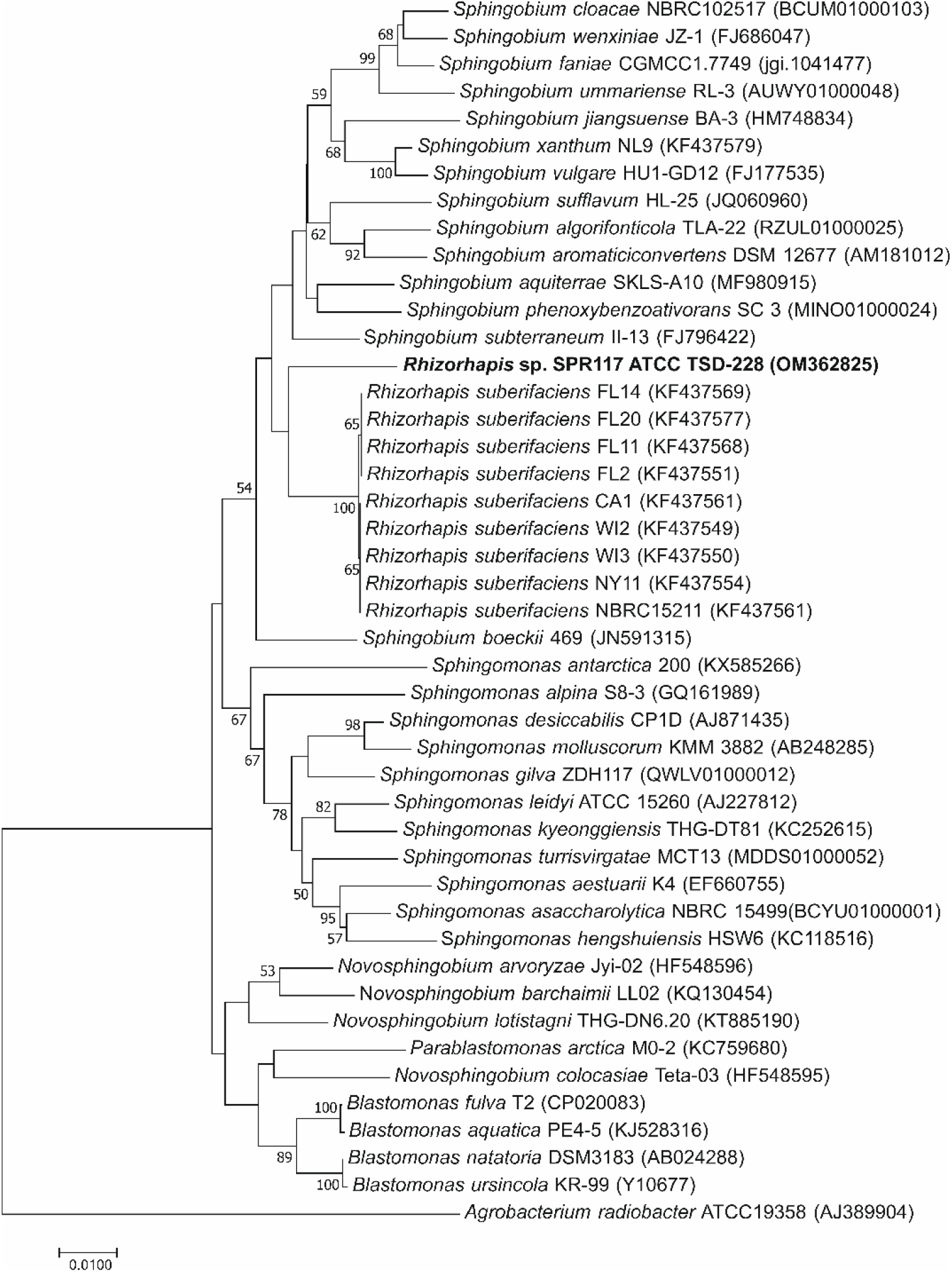
Neighbour-joining tree, based on 16S rRNA gene sequence data, showing the phylogenetic position of *Rhizorhapis* sp. strain SPR117 and most related microorganisms according to EzBioCloud. The genetic distances were calculated by using Kimura’s two-parameter model and the complete deletion option. Bootstrap values (calculated with 1000 resamplings) above 50% are shown in branch positions*. Agrobacterium radiobacter* ATCC19358 was used as outgroup organism. GenBank accession number of 16S rRNA gene sequences are given in parentheses. Bar, 0.01 expected changes per site.

### 3.3. *Characterization of* Rhizorhapis *sp. strain SPR117 (ATCC TSD-228)*

The genus *Rhizorhapis*, formerly known as *Rhizomonas*, belongs to the family *Sphingomonadaceae* (*Alphaproteobacteria* class) and currently comprises only the species type *Rhizorhapis suberifaciens,* which is causal agent of the lettuce (*Lactuca sativa* L.) corky root disease (Francis et al., 2014). Corky root has been one of the most important diseases of lettuce in production areas of USA, Europe, and Australia (van Bruggen et al., 2015). In severe cases of the disease, the entire taproot becomes brown, severely cracked, non-functional, and easily breakable (van Bruggen et al., 2014b). These symptoms at root level result in reduced head sizes of lettuce plants and decreasing crop yields. Corky root disease can be very severe —especially in intensive production systems and in warm climates—, with yield losses ranging from 30 to 70% (van Bruggen et al., 2014a). In this context, the evaluation of the phytopathogenic potential of the strain SPR117 was of interest.

We conducted a greenhouse experiment where the pathogenicity of SPR117 was tested in the lettuce varieties Romaine and Waldm’nn’s Dark Green. After 30 days, the root and shoot dry weight of lettuce plants was not affected by the inoculation of SPR117 (Fig. 4). The potential corky root severity in the inoculated plants was also evaluated according to the guidelines provided by Brown and Michelmore (1988), finding no symptoms related to the disease. Therefore, the strain SPR117 is not pathogenic to lettuce, at least to the varieties tested. Interestingly, some *Rhizorhapis* sp. strains have shown to provide specific control of corky root disease (van Bruggen et al., 2014b) and further studies will be needed in this respect for the strain SPR117.

**Fig. 4.**
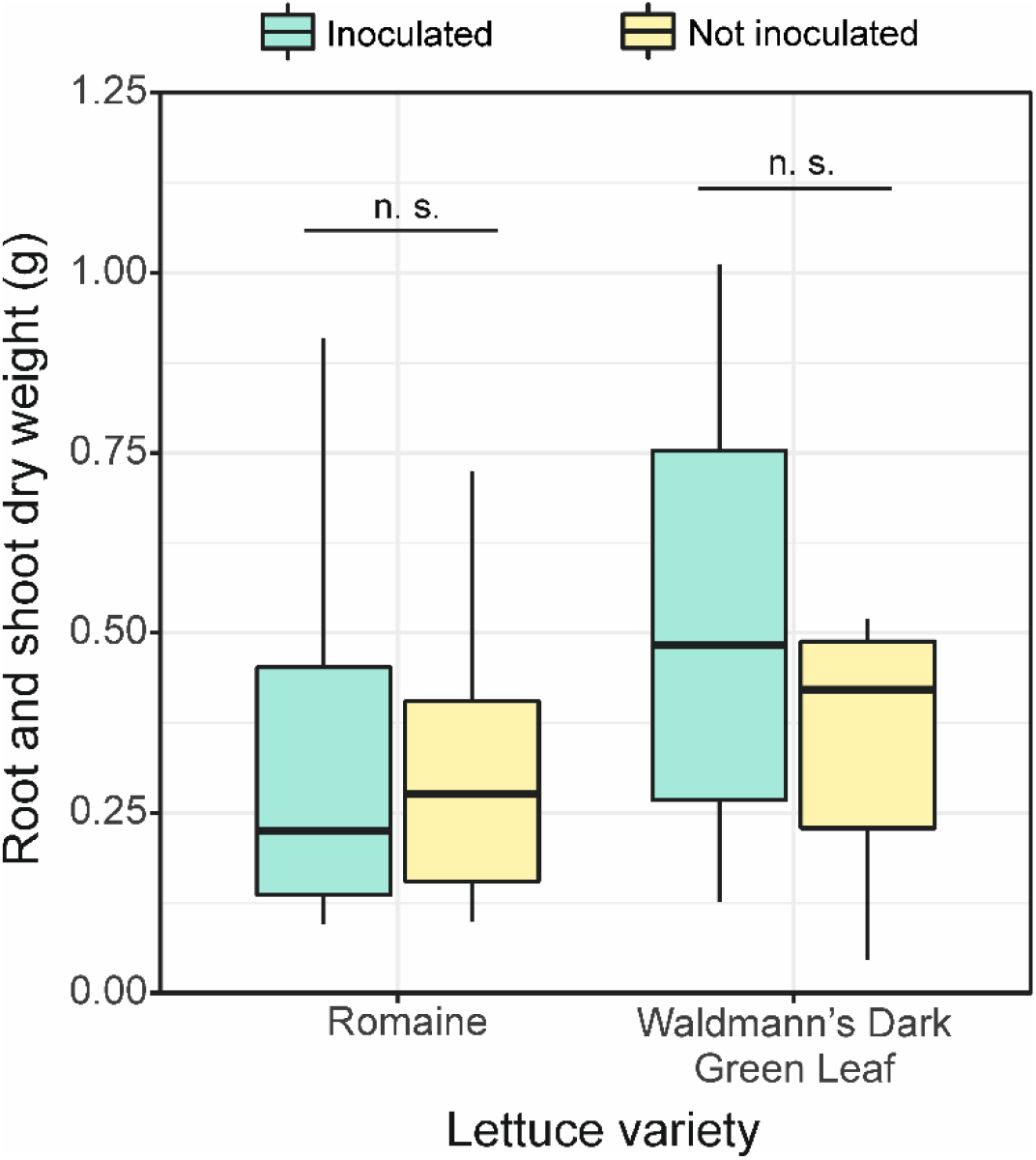
Box plots comparing root and shoot dry weight of lettuce plants of the varieties Romaine and Waldmann’s Dark Green Leaf after growing 28 days in sterilized potting substrate inoculated or not with *Rhizorhapis* sp. strain SPR117. The boxes represent the interquartile range (IQR) between the first and third quartiles (25^th^ and 75^th^ percentiles, respectively) and the vertical line inside the box defines the median. Whiskers represent the lowest and highest values within 1.5 times the IQR from the first and third quartiles, respectively. No significant differences were found between inoculated and non-inoculated plants at p<0.05 according to PERMANOVA.

Further insights into the strain SPR117 were obtained through the sequencing of its genome. Sequencing work yielded more than 33 M PE sequences, 18.79 M of which remained after quality-filtering and were subsequently used for assembling and annotating the genome. Main features of the assembly and annotation processes are summarized in Table 1. The genome of *Rhizorhapis* sp. SPR117 was assembled into 256 contigs (>300 bp), with a length of 4,419,522 bp and an N50 score of 62,062 bp. It has a GC content of 59.9%, similar to that of *R. suberifaciens* (58.9%) (Francis et al., 2014), and a completeness of 98%. The annotation process predicted a total of 4,278 coding genes and coding sequences (CDSs), 3 rRNAs, and 46 tRNAs (Table 1). 1129 CDSs were successfully classified into subsystems by RAST. A subsystem is a set of proteins that together implement a specific biological process or structural complex (Overbeek et al., 2005). The four most abundant subsystem categories were membrane transport, amino acids and derivatives, carbohydrates, and protein metabolism (Fig. S1).

**Table 1.** Assembly and annotation metrics of *Rhizorhapis* sp. strain SPR117

| Genome feature | *Rhizorhapis* sp. strain SPR117 |
| --- | --- |
| Sequence length (bp) | 4,419,522 |
| Contigs | 256 |
| Completeness (%) | 98.4 |
| Fine consistency (%) | 96.2 |
| GC content (%) | 59.86 |
| Longest contig length (bp) | 190,373 |
| N<sub>50</sub> | 62,062 |
| L<sub>50</sub> | 23 |
| Genes (total) | 4,452 |
| CDS<sup>1</sup> (total) | 4,399 |
| Genes (coding) | 4,278 |
| CDS<sup>1</sup> (coding) | 4,278 |
| rRNAs | 3 (1 5S, 1 16S, 1 23S) |
| tRNAs | 46 |
| ncRNAs | 4 |
| Pseudogenes | 121 |

<sup>1</sup>CDS: Coding sequences

Members of the genus *Rhizorhapis* have been hypothesized to degrade petroleum hydrocarbons (Brereton et al., 2020); however, the genome of the strain SPR117 does not present genes compatible with this capability. This is seen in concordance with the results of a laboratory experiment where we observed lack of bacterial growth when the isolate SPR117 was cultivated in culture media containing alkanes (dodecane) or PAHs (anthracene, benzene, and phenanthrene) as the only C source.

Metals, both those with and without a biological function, can be toxic for bacteria when accumulated to a certain concentration in the environment (Valls and de Lorenzo, 2002). Their intake by microorganisms is thus subjected to complex homeostatic mechanisms that ensure sufficient, but not excessive, acquisition (Chen et al., 2019; Valls and de Lorenzo, 2002). Furthermore, bacteria have developed mechanisms of metal resistance, which involve intra- or extracellular immobilization, biotransformation of the toxic metal ion into a less noxious form, dissimilatory reduction, and modulation of metal concentration homeostasis through cation efflux pumps (Romero et al., 2021). Long-term metal polluted ecosystems, as the waste sediment studied here, have been recognized as potential hotspots for the isolation of as-yet-uncultured bacteria with metal resistance capabilities (Abbas et al., 2015; Vasconcelos et al., 2021). In line with this argumentation, we found that the strain SPR117 has potential metal-tolerance capabilities since its genome harbors a variety of genes encoding proteins involved in the resistance to arsenic, cadmium, cobalt, copper, iron, mercury, and zinc, among others (Table S4). This finding is seen in connection with the results of the work by Brereton et al. (2020), which showed that the relative abundance of OTUs taxonomically affiliated to the genus *Rhizorhapis* increased in the rhizosphere of *Festuca arundinacea* in comparison with the unplanted treatment during the phytoremediation of different metal-contaminated soils. Among the aforementioned metals, the sediment presented high levels of arsenic, copper, iron, and mercury (Table S1); we thus dug deeper into the genes related to the resistance to these metals in the genome of *Rhizorhapis* sp. SPR117.

*Rhizorhapis* sp. SPR117’s genome contains genes related to arsenic methylation (*arsH* gene) and reduction (*arsC*, *arsO*, *arsR* and *acr3*) (Table S4). The *arsC* gene encodes a glutaredoxin arsenate reductase that reduces arsenate to arsenite within the cytoplasmic membrane; arsenite is then exported out of the cell by an efflux bum (Abdullahi et al., 2021; Li et al., 2021). In fact, the *acr3* gene encodes an arsenite efflux permease (Li et al., 2021). The presence of *arsH* in SPR117 may point out its capability to oxidize methyl arsenate compounds (Abbaszade et al., 2020). The four main pathways of microbial As detoxification in the environment are arsenite oxidation, arsenate reduction, arsenate respiration and arsenate methylation (Li et al., 2021).

Copper has an important role in cellular function. Due to its ability to undergo redox changes [Cu(I) ↔ Cu(II)], copper is an ideal cofactor for enzymes catalyzing electron transfers. However, this characteristic also makes copper dangerous for a cell as it can be involved in Fenton-like reactions creating reactive oxygen species. Cu(I) can attack and destroy iron-sulfur clusters, which releases iron that can in turn cause oxidative stress. Therefore, copper homeostasis has to be highly balanced to ensure proper cellular function and avoid cell damage (Rensing and McDevitt, 2013). Genes for copper resistance detected in the genome of the strain SPR117 included *copB*, *copC* and *copD* (Table S4). *copB* and *copD* encode outer and inner membrane proteins, respectively, while the CopC protein is believed to be a periplasmic copper chaperone (Lawton et al., 2016). Other genes for copper resistance were *cusA*, *cusB*, and *CusC*, which encode three proteins that are believed to form a complex that spans the periplasm and has the ability to remove copper and silver from the cell (Lawton et al., 2016).

The microbial iron homeostasis is highly regulated as this metal is required for cellular processes, but it becomes toxic at high concentrations (Abdullahi et al., 2021). Several genes responsible for iron metabolism and transport were found in the genome of the strain SPR117 (Table S4). Iron-sulfur enzymes (as those synthetized by the *sufA*, *sufB*, *sufC*, and *sufD* genes) are involved in important biological functional and processes such as electron transfer, regulating gene expression and substrate binding/activation, where the active site is made of iron and sulfur atoms (Abdullahi et al., 2021).

Mercuric ions are toxic to bacteria because they bind to sulfhydryl groups and hamper macromolecule synthesis and enzyme functioning (Abbaszade et al., 2020). Mercury resistance system of *Rhizorhapis* sp. SPR117 consists of the *merA*, *merP*, *merR,* and *merT* genes (Table S4). The reductase encoded by the *merA* gene is able to transform Hg^2+^ (highly reactive and toxic) into Hg^0^ (less toxic and volatile) (Arregui et al., 2021). *merP* and *merT* encode the proteins that conform the well-known MerP-MerT mercury transporter (Valls and de Lorenzo, 2002). Mercury resistance genes are usually organized in an operon, which is repressed or activated by the product of the *merR* gene. Our results thus point towards the potential high versatility, and unknown so far, of members of the genus *Rhizorhapis* to deal with metal contamination.

Although metals are mostly not susceptible to complete microbial degradation, bioremediation of metal pollution uses microorganisms to immobilize or transform metals into non-bioavailable or less toxic forms (Ye et al., 2017). These approaches do not solve the problem completely, but they do help to protect polluted sites from noxious effects and isolate the contaminants as a contained, and sometimes, recyclable unit (Valls and de Lorenzo, 2002). Newly isolated bacteria, such as *Rhizorhapis* sp. strain SPR117, with metal tolerant features are a potential source of bioremediation tools. For example, as shown by Teixeira et al. (2014) and Emenike et al. (2015), bioaugmentation of metal-contaminated soils has shown promising results. On the other hand, metal resistance genes are valuable resources for genetic engineering. For instance, the transformation of *Populus alba* × *P. tremula* var. *glandulosa* with a metal tolerance gene (*ScYCF1*, yeast cadmium factor 1) led to improved poplar growth and cadmium accumulation in aerial tissue (Shim et al., 2013). The overexpression of a phytochelatin synthase from *Schizosaccharomyces pombe* in *Escherichia coli* resulted in a higher cellular cadmium accumulation than that in the wild type (Kang et al., 2007). And the engineering of *Pseudomonas putida* for expression of the *arsM* arsenic(III) S-adenosine methyltransferase gene showed to be an effective approach to volatilize arsenic from polluted soils (Chen et al., 2014). In line with these examples, *Rhizorhapis* sp. SPR117 could be used to (i) bioremediate metal-polluted soils or sediments through bioaugmentation (once its metal bioremediation capabilities are experimentally confirmed) and/or (ii) develop bio-engineered tools involving its metal resistance genes.

## 4. Conclusions

In the present work, we coupled 16S rRNA gene metabarcoding and plate culturing (with further colony picking and identification of the isolates) to taxonomically describe the bacterial community inhabiting multi-contaminated (petroleum hydrocarbons and heavy metals) waste sediment collected from a former industrial dumpsite. The bacterial diversities recovered by both approaches greatly differed and only a small fraction (2.7%) of the culture-independent bacterial community was cultured due to the current incapability to reproduce in the laboratory the specific abiotic and biotic conditions that most bacteria need to growth. Surprisingly, most of the culturable OTUs either were absent or found in very low abundances in the culture-independent community, which demonstrates that culturing complements the diversity retrieved through metataxonomics and is a useful tool to investigate the rare bacterial biosphere. In fact, one member of this rare biosphere in the studied waste sediment was identified as a potential new species in the genus *Rhizorhapis*. The genome sequence analysis of *Rhizorhapis* sp. strain SPR117 (ATCC TSD-228) demonstrated that this strain may have bioremediation capabilities since genes related to metal resistance were annotated. This shows that rare biosphere is a reservoir of potential tools for bioremediation. Further studies will include a formal description of *Rhizorhapis* sp. strain SPR117 as a new species and will assess the metal bioremediation capabilities of this bacterium under real conditions.

## CRediT authorship contribution statement

**José A. Siles:** Conceptualization, Methodology, Formal analysis, Investigation, Writing - Original Draft, Visualization. **Andrew J. Hendrickson:** Investigation, Writing - Review & Editing. **Norman Terry**: Resources, Funding acquisition, Writing - Review & Editing.

## Acknowledgements

The authors thank Ben LePage and Bob Grey from PG&E Company for providing us access to Shell Pond and Katherine French for her comments on an earlier draft of the manuscript.

## Data availability

Sequencing data have been deposited in public repositories. Other used data will be made available on request.

