## Supplementary Material for "Coupling of metataxonomics and culturing improves bacterial diversity characterization and identifies a novel *Rhizorhapis* sp. with metal resistance potential in a multi-contaminated waste sediment"

***for***

**The supplementary material includes Supplemental Tables S1-S4 and Figure S1**

Table S1. Main physicochemical properties and levels of contaminants in the studied waste sediment.

| | Average ± Standard deviation |
|---|---|
| **Texture** | |
| Sand (%) | 38.67 ± 0.58 |
| Silt (%) | 21.67 ± 0.58 |
| Clay (%) | 39.67 ± 0.58 |
| **Chemical properties** | |
| pH | 8.23 ± 0.11 |
| Electrical conductivity (mS $cm^{-1}$) | 1.19 ± 0.06 |
| Total carbon (mg $g^{-1}$) | 528 ± 25 |
| Organic carbon (mg $g^{-1}$) | 524 ± 4 |
| Total nitrogen (mg $g^{-1}$) | 2.63 ± 0.04 |
| C/N | 199 ± 4.5 |
| Total phosphorus (mg $g^{-1}$) | 7.2 ± 1.72 |
| **Petroleum hydrocarbons** | |
| Total petroleum hidrocarbons ($C_8$-$C_{40}$) (g $kg^{-1}$) | 20.14 ± 2.16 |
| Total PAHs (mg $kg^{-1}$) | 370 ± 89 |
| **PAHs** | |
| Benzo(*ghi*)perylene (mg $kg^{-1}$) | 190 ± 26 |
| Pyrene (mg $kg^{-1}$) | 74.0 ± 31.4 |
| Indeno(1,2,3-*cd*)pyrene (mg $kg^{-1}$) | 21.3 ± 1.5 |
| Benzo(*a*)pyrene (mg $kg^{-1}$) | 20.0 ± 2.7 |
| Fluoranthene (mg $kg^{-1}$) | 19.3 ± 3.3 |
| Phenanthrene (mg $kg^{-1}$) | 14.5 ± 2.5 |
| Acenaphthylene (mg $kg^{-1}$) | 12.3 ± 2.3 |
| Benzo(*b*)fluoranthene (mg $kg^{-1}$) | 6.2 ± 1.2 |
| Benzo(*k*)fluoranthene (mg $kg^{-1}$) | 2.2 ± 0.2 |
| Benzo(*a*)anthracene (mg $kg^{-1}$) | 1.9 ± 0.2 |
| Others (mg $kg^{-1}$) | 9.2 ± 2.3 |
| **Heavy metals** | |
| Chromium (mg $kg^{-1}$) | 220 ± 12.3 |
| Barium (mg $kg^{-1}$) | 160 ± 15.1 |
| Zinc (mg $kg^{-1}$) | 150 ± 9.8 |
| Copper (mg $kg^{-1}$) | 120 ± 11.2 |
| Lead (mg $kg^{-1}$) | 88 ± 4.3 |
| Nickel (mg $kg^{-1}$) | 78 ± 2.3 |
| Vanadium (mg $kg^{-1}$) | 42 ± 4.6 |
| Molybdenum (mg $kg^{-1}$) | 65 ± 3.1 |
| Cobalt (mg $kg^{-1}$) | 16 ± 0.9 |
| Arsenic (mg $kg^{-1}$) | 16 ± 1.1 |
| Mercury (mg $kg^{-1}$) | 3.1 ± 0.9 |

Table S2. Relative abundance and taxonomic affiliation of OTUs in the culture-independent community with a relative abundance >1%.

| Otu ID | Phylum | Identity (%) | Class | Identity (%) | Genus | Identity (%) | Relative abundance (%) |
|---|---|---|---|---|---|---|---|
| Otu1 | *Bacteroidetes* | 100 | *Flavobacteriia* | 100 | *Hoppeia* | 59 | 46.77 |
| Otu2 | *Proteobacteria* | 100 | *Gammaproteobacteria* | 100 | *Kangiella* | 100 | 3.91 |
| Otu12 | *Proteobacteria* | 59 | *Deltaproteobacteria* | 54 | *Desulfohalophilus* | 14 | 3.27 |
| Otu5 | *Saccharibacteria* | 100 | | | | | 3.18 |
| Otu7 | *Verrucomicrobia* | 100 | *Opitutae* | 100 | *Pelagicoccus* | 84 | 2.37 |
| Otu3 | *Bacteroidetes* | 100 | *Flavobacteriia* | 88 | *Owenweeksia* | 44 | 2.19 |
| Otu4 | *Balneolaeota* | 100 | *Balneolia* | 100 | *Gracilimonas* | 100 | 2.03 |
| Otu6 | *Actinobacteria* | 100 | *Acidimicrobiia* | 91 | *Iamia* | 42 | 1.88 |
| Otu9 | *Proteobacteria* | 100 | *Gammaproteobacteria* | 100 | *Pseudidiomarina* | 100 | 1.34 |
| Otu10 | *Chloroflexi* | 64 | *Thermomicrobia* | 57 | *Nitrolancea* | 26 | 1.30 |
| Otu8 | *Proteobacteria* | 100 | *Gammaproteobacteria* | 100 | *Pseudidiomarina* | 100 | 1.08 |

Table S3. Taxonomic information and relative abundance of the culturable OTUs and their phylogenetic relationship with the culture-independent OTUs. cOTU = Culturable OTU. mOTU = metabarcoding OTU. RA = Relative abundance.

| cOTU ID | Isolates OTU | cOTU RA (%) | Isolate ID | Closest relative Eztaxon match (GenBank accession no.), %similarity of the OTU representative isolate | Phylum | Related mOTU | RA (%) of the mOTU |
|---|---|---|---|---|---|---|---|
| Otu1 | 39 | 27.86 | SPR114 | *Bacillus thioparans* BMP-1 (DQ371431), 100 | *Firmicutes* | Otu440 | 0.05 |
| Otu2 | 17 | 12.14 | SPL145 | *Bacillus oryzaecorticis* R1 (KF548480), 99.11 | *Firmicutes* | Otu287 | 0.03 |
| Otu3 | 13 | 9.29 | SPR106 | *Fictibacillus phosphorivorans* Ca7T (JX258924), 99.70 | *Firmicutes* | | |
| Otu4 | 7 | 5.00 | SPR136 | *Demequina activiva* BS-12M (KM591918), 99.35 | *Actinobacteria* | Otu88 | 0.07 |
| Otu5 | 6 | 4.29 | SPR6 | *Pseudomonas alcaligenes* NBRC 14159 (BATI01000076), 98.97 | *Proteobacteria* | Otu36 | 0.23 |
| Otu6 | 9 | 6.43 | SPR140 | *Caenimicrobium hargitense* CGII-59m2 (KM083134), 97.81 | *Proteobacteria* | Otu307 | 0.01 |
| Otu7 | 4 | 2.86 | SPL106 | *Lysobacter arseniciresistens* ZS79 (AVPT01000055), 97.27 | *Proteobacteria* | Otu26 | 0.39 |
| Otu8 | 4 | 2.86 | SPL34 | *Pseudomonas kunmingensis* HL22-2 (JQ246444), 99.14 | *Proteobacteria* | Otu93 | 0.28 |
| Otu9 | 5 | 3.57 | SPR46 | *Microbacterium oxydans* DSM 20578 (Y17227), 99.82 | *Actinobacteria* | | |
| Otu10 | 2 | 1.43 | SPR17 | *Bacillus zhangzhouensis* DW5-4 (JOTP01000061), 99.91 | *Firmicutes* | Otu543 | 0.01 |
| Otu11 | 1 | 0.71 | SPL5 | *Planococcus citreus* DSM 20549 (RCCP01000013), 100 | *Firmicutes* | Otu580 | 0.02 |
| Otu12 | 1 | 0.71 | SPL154 | *Streptomyces yanii* NBRC 14669 (AB006159), 100 | *Actinobacteria* | Otu57 | 0.15 |
| Otu13 | 1 | 0.71 | SPL27 | *Bacillus firmus* NBRC 15306 (BCUY01000205), 99.40 | *Firmicutes* | Otu360 | 0.05 |
| Otu14 | 2 | 1.43 | SPR19 | *Bacillus onubensis* 0911MAR22V3 (NSEB01000017), 100 | *Firmicutes* | | |
| Otu15 | 2 | 1.43 | SPR125 | *Mesorhizobium sediminum* YIM M12096 (KX151664), 99.72 | *Proteobacteria* | | |
| Otu16 | 2 | 1.43 | SPR132 | *Georgenia muralis* DSM 14418 (RKRA01000001), 99.33 | *Actinobacteria* | | |
| Otu17 | 1 | 0.71 | SPL20 | *Brevibacterium sandarakinum* DSM 22082 (LT629739), 97.03 | *Actinobacteria* | Otu202 | 0.01 |
| Otu18 | 1 | 0.71 | SPR144 | *Cellulomonas carbonis* T26 (HQ702749), 99.25 | *Actinobacteria* | Otu410 | 0.01 |
| Otu19 | 2 | 1.43 | SPL19 | *Staphylococcus saprophyticus* ATCC 15305 (AP008934), 100 | *Firmicutes* | | |
| Otu20 | 2 | 1.43 | SPL22 | *Pseudomonas oleovorans* subsp. *lubricantis* RS1 (DQ842018), 98.45 | *Proteobacteria* | | |
| Otu21 | 2 | 1.43 | SPR21 | *Agrobacterium salinitolerans* YIC 5082 (MRDH01000011), 100 | *Proteobacteria* | | |
| Otu22 | 1 | 0.71 | SPR141 | *Nocardioides massiliensis* GD13 (CCXJ01000100), 97.29 | *Actinobacteria* | Otu172 | 0.02 |
| Otu23 | 3 | 2.14 | SPR42 | *Actinotalea ferrariae* CF5-4 (HQ730135), 98.03 | *Actinobacteria* | | |
| Otu24 | 1 | 0.71 | SPL24 | *Bosea robiniae* DSM 26672 (jgi.1085745), 99.18 | *Proteobacteria* | | |
| Otu25 | 1 | 0.71 | SPL31 | *Photobacterium halotolerans* MACL01 (AY551089), 97.65 | *Proteobacteria* | | |
| Otu26 | 1 | 0.71 | SPL32 | *Rheinheimera aquimaris* SW-353 (EF076757), 99.23 | *Proteobacteria* | | |
| Otu27 | 1 | 0.71 | SPL37 | *Vibrio japonicus* JCM 31412 (LC143378), 99.09 | *Proteobacteria* | | |
| Otu28 | 1 | 0.71 | SPR109 | *Sphingomonas laterariae* LNB2 (jgi.1118286), 97.66 | *Proteobacteria* | | |
| Otu29 | 1 | 0.71 | SPR115 | *Sphingopyxis alaskensis* RB2256 (CP000356), 99.83 | *Proteobacteria* | | |
| Otu30 | 1 | 0.71 | SPR116 | *Sphingomonas rubra* CGMCC 1.9113 (jgi.1058074), 97.68 | *Proteobacteria* | | |
| Otu31 | 1 | 0.71 | SPR117 | *Rhizorhapis suberifaciens* CA1 (KF437561), 96.88 | *Proteobacteria* | | |
| Otu32 | 1 | 0.71 | SPR13 | *Methylorubrum salsuginis* MR (EF015478), 99.19 | *Proteobacteria* | | |
| Otu33 | 1 | 0.71 | SPR130 | *Paenibacillus typhae* CGMCC 1.11012 (jgi.1076256), 98.34 | *Firmicutes* | | |
| Otu34 | 1 | 0.71 | SPR131 | *Bacillus niacini* IFO 15566 (AB021194), 99.51 | *Firmicutes* | | |
| Otu35 | 2 | 1.43 | SPR126 | *Cellulosimicrobium cellulans* LMG 16121 (CAOI01000359), 100 | *Actinobacteria* | | |

Table S4. Metal-resistance features in the genome of *Rhizorhapis* sp. strain SPR177.

| Heavy metal | Gene | Description |
|---|---|---|
| **Arsenic** | | |
| | *arsH* | Arsenic resistance protein ArsH |
| | *acr3* | Arsenical-resistance protein ACR3 |
| | *arsO* | Flavin-dependent monooxygenase ArsO associated with arsenic resistance |
| | *arsR* | Arsenical resistance operon repressor |
| | *arsC* | Arsenate reductase (EC 1.20.4.1) glutaredoxin-coupled, glutaredoxin-like family |
| | *arsC* | Arsenate reductase (EC 1.20.4.4) thioredoxin-coupled, LMWP family |
| **Cobalt** | | |
| | *corC* | Cobalt and magnesium efflux protein |
| **Copper** | | |
| | *copA* | Copper-exporting P-type ATPase |
| | *copB* | Copper resistance protein B |
| | *copC* | Copper resistance protein CopC |
| | *copD* | Copper resistance protein CopD |
| | *cusA* | Copper/silver efflux RND transporter, transmembrane protein |
| | *cusB* | Copper/silver efflux RND transporter, membrane fusion protein |
| | *cusC* | Copper/silver efflux RND transporter, outer membrane protein |
| | *cueO* | Multicopper oxidase |
| **Mercury** | | |
| | *merA* | Mercuric reductase |
| | *merP* | Periplasmic mercury (+2) binding protein |
| | *merR* | Mercuric resistance operon regulatory protein |
| | *merT* | Mercuric transport protein MerT |
| **Iron** | | |
| | *fhuA* | Ferrichrome-iron receptor |
| | *feoA* | Ferrous iron transporter A |
| | *feoB* | Ferrous iron transporter B |
| | *fieF* | Ferrous-iron efflux pump |
| | *sufA* | Iron-sulfur cluster assembly iron binding protein SufA |
| | *sufB* | Iron-sulfur cluster assembly protein SufB |
| | *sufC* | Iron-sulfur cluster assembly ATPase protein SufC |
| | *sufD* | Iron-sulfur cluster assembly protein SufD |
| | *iscR* | Iron-sulfur cluster regulator IscR |
| | *apbC* | Scaffold protein for [4Fe-4S] cluster assembly ApbC, MRP-like |
| | NK55_07155 | Iron binding protein from the HesB_IscA_SufA family |
| | | Putative iron-regulated membrane protein |
| | CZ765_12365 | Uncharacterized iron-regulated membrane protein; Iron-uptake factor PiuB |
| **Miscellaneous** | | |
| | czcD | Cobalt/zinc/cadmium resistance protein |
| | zntA/copA | Lead, cadmium, zinc and mercury transporting ATPase; Copper-translocating P-type ATPase |
| | | RcnR-like protein clustered with cobalt-zinc-cadmium resistance protein CzcD |

Fig. S1. Overview of subsystems for *Rhizorhapis* sp. strain SPR117.

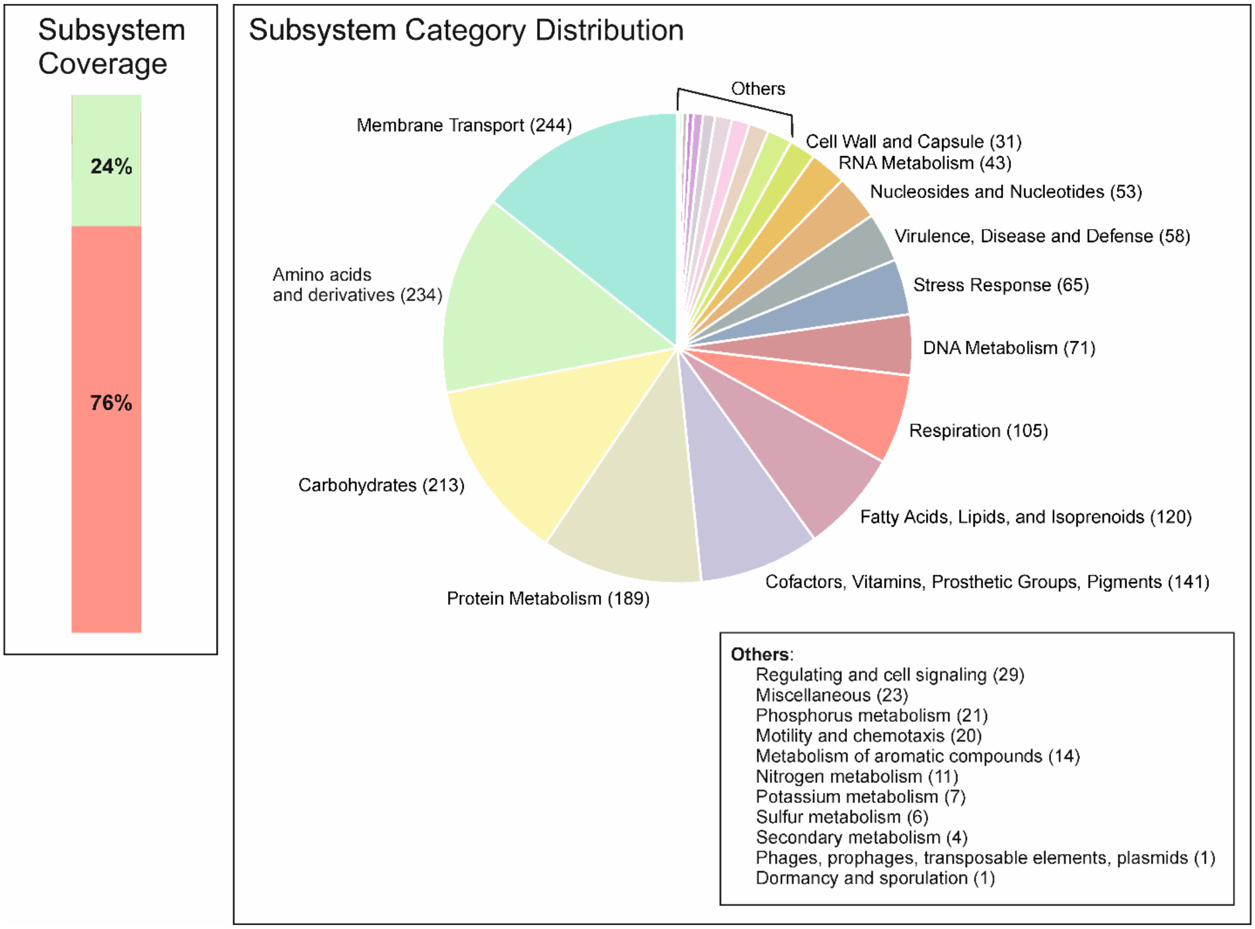
